# Diverse strategies link growth rate and competitive ability in phytoplankton responses to changes in CO_2_ levels

**DOI:** 10.1101/651471

**Authors:** Sinead Collins, C. Elisa Schaum

## Abstract

Aquatic microbial primary producers exist in genetically variable populations, but are often studied as single lineages. However, the properties of lineages grown alone often fail to predict the composition of microbial assemblages. We demonstrate that different lineages of a marine picoplankter have unique growth strategies, and that they modulate their lineage growth rate in the presence of other lineages. This explains why growth rates of lineages in isolation do not reliably predict the lineage composition of assemblages. The diversity of growth strategies observed are consistent with lineage-specific energy-allocation that depends on social milieu. Since lineage growth is only one of many traits determining fitness in natural assemblages, we propose that in all but the poorest quality environments where allocating maximum energy to growth is the only viable strategy, we should expect intraspecific variation in growth strategies, with more strategies possible in ameliorated environments, such as high CO_2_ for many marine picoplankton. This emphasizes the importance of understanding and accounting for basic organismal biology in our models of aquatic primary producers.

**Data archiving:** Should this manuscript be accepted, data will be archived at Pangea, and the DOI be included at the end of the article

---

Microbial primary producers form the base of aquatic ecosystems, and link organisms to their environment through their role in nutrient cycling^1,2^. Currently, populations of aquatic primary producers are usually modelled scaling up from growth rates measured in mono-cultures ^3^. This is important because the lineage composition of a population in turn affects bulk population properties, such as primary production or nutrient uptake, when there is also intraspecific variation in individual-level traits that underlie population properties. However, recent studies show that lineage growth in monoculture can be poor predictors of the composition of multilineage assemblages ^4-6^, highlighting that a more nuanced understanding of the impact of intraspecific diversity on trait values (for example on lineage growth or photosynthetic rates) is needed. Given the high intraspecific diversity in cell division rates in single celled primary producers ^7-11^, the lineage composition of a species has the potential to affect bulk traits of populations such as primary production, nutrient uptake, and trophic energy transfer.

Our understanding of the lineage frequencies in a population of closely-related lineages (such as conspecifics or lineages within a species complex) is based on competition theory where relative lineage growth rates are determined by differences in the uptake and metabolism of nutrients, often modulated by temperature and light ^12^. Here, we define a lineage to be cells related by descent with little enough genetic variation introduced by mutation over the timescale considered, that individual cells of the lineage have the same phenotype under the same environmental and social conditions. Major genetic changes to a focal lineage during reproduction, such as recombination events or horizontal gene transfer, are likely to result in a new lineage, whereas asexual reproduction of a focal lineage over timescales considered here are unlikely to result in a new lineage.

While competition theory does explain how differences in nutrient, temperature and light related traits (hereafter “metabolic traits”) affect relative fitness under different environmental conditions, metabolic trait values of a lineage are only determined by lineage identity and the abiotic environment. In this case, non-self conspecifics affect the population growth rate of a focal lineage through their effect on the abiotic environment, for example, by sequestering resources^13^. Here, relative lineage growth rates from single-lineage cultures (hereafter “monoculture”) would be good predictors of lineage frequencies where multiple lineages are grown together (hereafter “mixed culture”) in that same environment and population density.

Despite these clear predictions, monoculture growth rates can fail to correlate positively with competitive outcomes ^4,5^, in high-nutrient, low-cell density environments where rapid lineage growth should be favoured, and where results cannot be plausibly explained by higher-affinity, slower resource acquisition or use being advantageous. Importantly, the outcomes of competitions are repeatable, even when competing lineages have similar growth rates in monoculture under the same environmental conditions and cell densities ^4,6^. These data suggest that lineages modulate lineage growth rates in response to their social milieu, even if there is no change in the abiotic environment.

Additionally, there is high intraspecific or intra-population variation in cell division rate responses to environmental change^7-11^ and models suggest that intraspecific variation in cell division rates may be adaptive even at the level of single populations ^14^. Microbes have a wide range of strategies for lineage growth in diverse populations ^15^, from straightforward competition for a limiting resource to interdependence based on public goods ^16^. The role of variation in these growth strategies (defined here as the allocation of energy to different components of fitness, including, but not limited to lineage growth, production of resting stages, defence, and nutrient storage) in determining the biogeochemical role of phytoplankton in different environments is established ^17^. Intraspecific variation in growth strategies in the same abiotic environment, and the ability of individual lineages to modulate growth strategies in response to changes in species diversity are less studied, but vitally important for scaling up from single to multi-lineage populations.

Here, we explore variation in the relationship between the population growth rates of focal lineages in monoculture and mixed culture at ambient and elevated pCO_2_. First, we show that *Ostrecococcus* detects and react to non-self conspecifics, such that lineage growth rates in monoculture and mixed culture can differ. Second, we argue that multiple growth strategies can co-exist in populations of closely related lineages and that there is thus potential for variation in the relationship between lineage growth rates and social milieu. We show that this variation is higher under ameliorated conditions, such as carbon enrichment. This variation in lineage responses and growth strategies explains why single-lineage experiments often fail to predict the composition and bulk properties of assemblages, and reconciles these seemingly disparate experiments with a single overarching explanation.

## RESULTS and DISCUSSION

### Multiple growth-competition strategies are observed in laboratory experiments

Cell division rate is only one component of fitness in microbes, and the organisms that we bring into the laboratory have long evolutionary histories where cell division rate probably was not the sole determinant of overall fitness. Here, we experimentally tested whether microbial primary producers respond differently to self and non-self conspecifics in the absence of nutrient limitation or obvious antagonistic interactions, and whether these responses are lineage-specific strategies that modulate lineage growth rate. We exposed at least 6 lineages of *Ostreococcus sp.* to three different non-self cues using self cues and fresh 0.2µm filtered culture medium as a control (see Fig. 1 A-D for a schematic of the experiment) 1) non-self live cells directly, 2) non-self cells on the other side of a permeable barrier or 3) non-self, nutrient-supplemented, spent, cell-free media. All tests were carried out in nutrient-replete media at low cell densities (100 cells mL^-1^ to 10^5^ cells mL^-1^). To date, no quorum-sensing or antagonistic (i.e. – toxin producing) behaviours have been observed in *Ostreococcus*.

**Figure 1.**
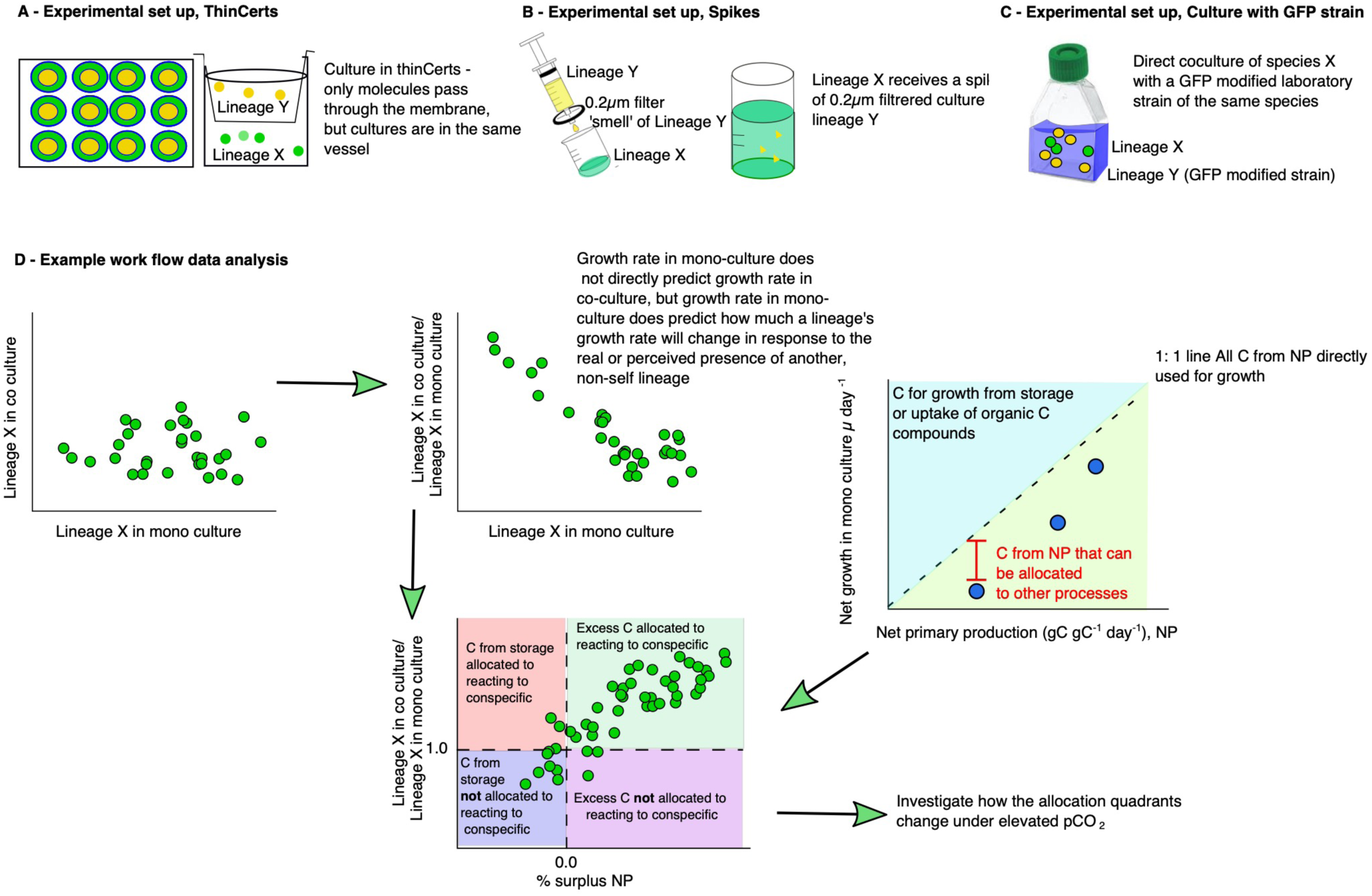
Schematic overview of work flow. For this experiment, we used ambient and elevated pCO_2_ evolved *Ostreococcus* lineages after 400 generations of selection. First, we tested their ability to react to the direct, indirect, or perceived presence of conspecifics. A) In the ‘indirect’ presence scenario, samples were cultured in wells divided by a semi-permeable membrane (ThinCert). B) For the ‘perceived’ presence scenario, we used 0.2µm filtrate of an exponentially growing sample (hereafter referred to as ‘Spike’). C) For the ‘direct’ presence scenario samples were grown alongside an GFP-transformed *Ostreococcus* lineage. D) Overview of analysis. We first compared growth rates of a lineage X in mono-culture to that same lineage in co-culture. In a next step, we calculated the *ratio* of growth in co-culture vs growth in mono-culture, to get a measure of how much growth rates *change* in response to the (direct, indirect, or perceived) presence of another lineage from the same species complex. For the lineages in mono-culture, we also measured net photosynthesis rates at their selection pCO_2_. Converting growth rates and photosynthesis rates into units carbon allows us to determine whether lineages must use storage C-sources to achieve the growth rates determined here or produce excess photosynthate, which can be exuded, stored, or used for processes other than growth (note that NP contains ‘losses’ from respiration^41^). Finally, we can form hypotheses about how this excess carbon could be used in reactions to conspecifics by testing the relationship between excess carbon and reactions to conspecifics over all lineages. This does not describe the causal underlying molecular mechanism, but yields highly repeatable results that predict lineage reactions to conspecifics.

Regardless of the method of exposure, lineages evolved under ambient (Fig. 2 A-C, 400ppm CO_2_) or elevated pCO_2_ (Fig. 3 A-C, 1000ppm CO_2_) changed their cell division rates in response to non-self signals (for statistics, see supplementary Tables 1-2 and 5-6 for statistical models on ambient and elevated pCO_2_ selected lineages respectively). The responsiveness of lineages to non-self cues was graded: direct co-culture with the GFP modified strain elicited the strongest response, followed by co-culture in a thin-cert, and finally, ‘spiked’ culture medium. Responses were repeatable *within* lineages but differed *between* lineages, demonstrating that responses are lineage-specific strategies that modulate lineage growth rate based on social cues. Thus, there is intraspecific variation in growth modulation in response to the presence to non-self conspecifics, even in a stable laboratory environment and without obvious competition for nutrients or other resources. In general, lineages that grow slowly in monoculture increase their growth rate in indirect co-culture or with supernatant spikes, while lineages that grow faster in monoculture either decrease or do not change their growth rate in response to the same stimuli (see Fig. 2 A and B, Fig. 3 A and B). When assayed in direct co-culture with a GFP modified strain, all ambient pCO_2_ evolved lineages (Fig. 2C), regardless of growth in monoculture, increase lineage growth rate, with slower growing lineages having a higher relative increase in growth than faster growing ones. These data demonstrate first, that *Ostreococcus* responds to signals from conspecifics by modulating lineage growth rate even in the absence of any change in the abiotic environment and second, that there is intraspecific variation in both the direction and magnitude of growth modulation caused by non-self cues. This is in line with previous studies showing different lineage growth rates in monoculture or in mixed culture ^18^.

**Figure 2.**
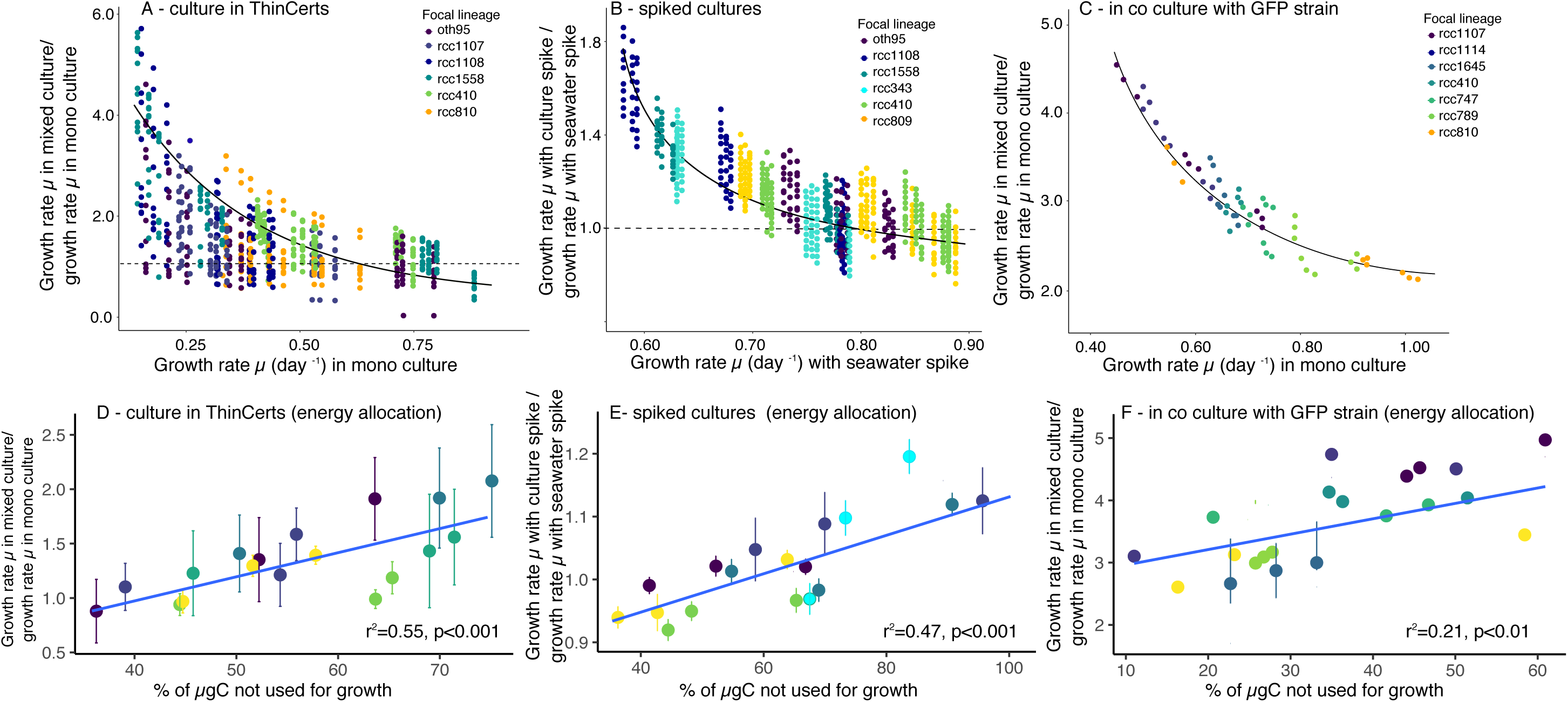
Ambient pCO_2_ selected lineages: Fold change of growth rate in mixed culture relative to growth rate in monoculture as a function of growth rate in mono culture for. A) lineages cultured in ThinCerts, B) lineages spiked with supernatant of conspecifics, C) lineages in mixed culture with a GFP *Ostreococcus* strain. In all cases, there is a trend for samples with high growth rates in mono-culture to have lower growth rates in mixed culture and vice versa. The fitted black lines in A-C are parameterised using non-linear mixed model output (supporting information Tables 1 and 2). The dashed line in panels A and B indicates a fold-change of 1. Values >1 indicate faster growth in mixed culture than in monoculture. **Fold-change in growth rates (as in A-C) as a function of photosynthetic carbon allocation to biomass in cultures grown in D) indirect co-culture, E) spiked non-self media and F) co-culture with a GFP-transformed *Ostreococcus* strain**. % values indicate how much photosynthetically-fixed carbon is available to processes other than growth relative to the amount of photosynthetically-fixed carbon allocated to increase in biomass (i.e. a value of 50% indicates that half as much carbon as is put into growth can be made available for other processes). Values <0 indicate that lineages must be using internal storages or draw organic carbon from elsewhere. Lineages that allocate less carbon to biomass production increase their growth more in response to signals from non-self (see Supporting Tables 3 and 4 for statistics). Colours indicate focal lineage, error-bars indicate ± 1S.E.M. The fitted blue line is the output of a linear mixed effects model. For each unique focal lineage – non-self pair, n =3 (three biological replicates, with three technical replicates, and all lineages in full interaction with each other).

**Figure 3.**
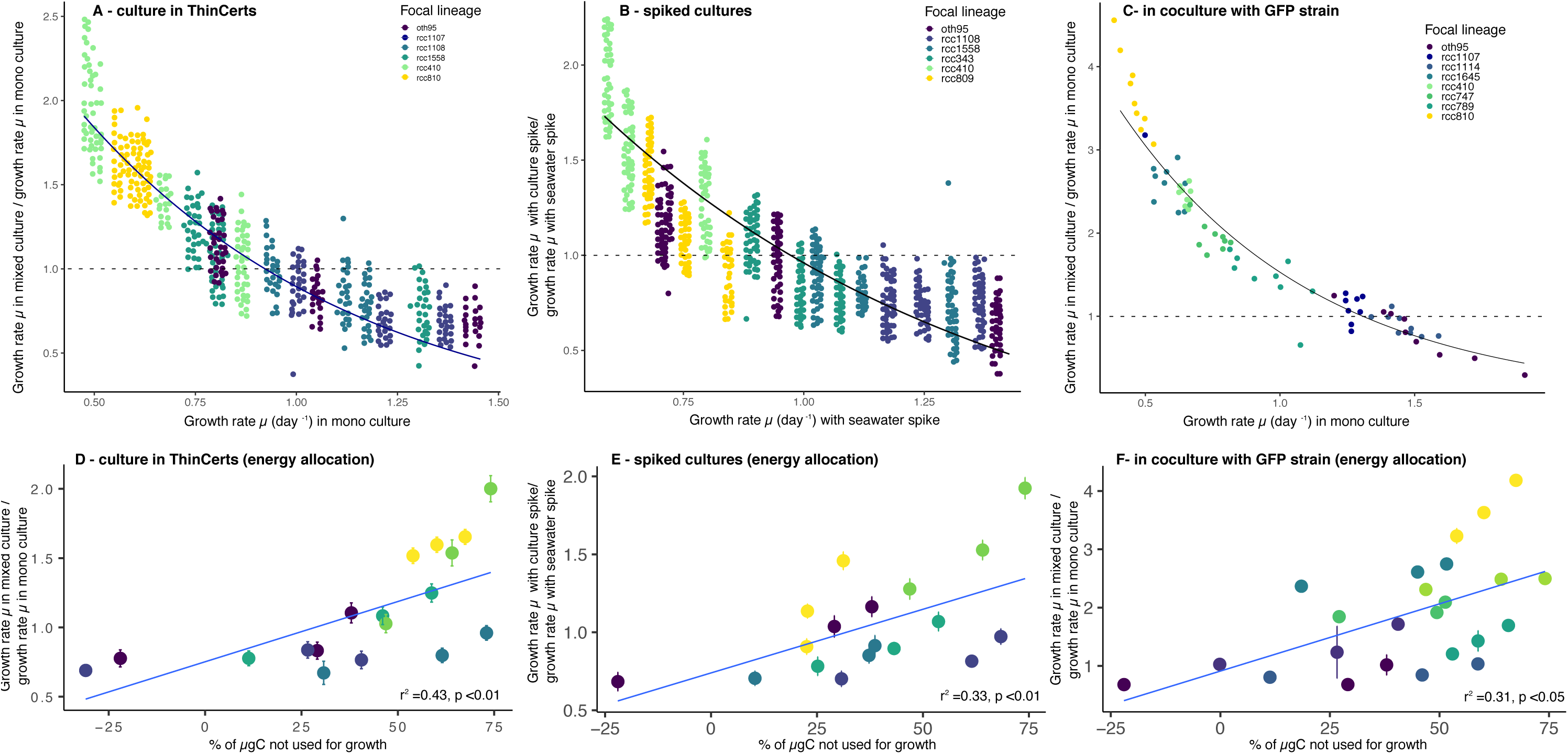
Elevated pCO_2_ selected lineages: Fold change of growth rate in mixed culture relative to growth rate in monoculture as a function of growth rate in mono culture for. A) lineages cultured in ThinCerts, B) lineages spiked with supernatant of conspecifics, C) lineages in mixed culture with a GFP *Ostreococcus* strain. In all cases, there is a trend for samples with high growth rates in mono-culture to have lower growth rates in mixed culture and vice versa. The fitted black lines in A-C are parameterised using non-linear mixed model output (supporting information Tables 5 and 6). The dashed line in panels A and B indicates a fold-change of 1. Values >1 indicate faster growth in mixed culture than in monoculture. Relative to ambient pCO_2_ evolved lineages, between-lineages are larger under elevated pCO_2_,while fold changes in growth are smaller (possibly because lineages are already growing faster under elevated pCO_2_), and even under direct co-culture, not all lineages increase their growth rates. **Fold-change in growth rates (as in A-C) as a function of photosynthetic carbon allocation to biomass in cultures grown in D) indirect co-culture, E) spiked non-self media and F) co-culture with a GFP-transformed *Ostreococcus* strain**. % values indicate how much photosynthetically-fixed carbon is available to processes other than growth relative to the amount of photosynthetically-fixed carbon allocated to increase in biomass (i.e. a value of 50% indicates that half as much carbon as is put into growth can be made available for other processes). Values <0 indicate that lineages must be using internal storages or draw organic carbon from elsewhere. Lineages that allocate less carbon to biomass production increase their growth more in response to signals from non-self (see Supporting Tables 7 and 8 for statistics). Relative to ambient pCO_2_ evolved samples, lineages from the elevated pCO_2_ treatment are more likely to follow different carbon allocation strategies, such as *not* allocating excess carbon to growth in the presence of a conspecific, or using storage carbon rather than excess carbon to react to conspecifics. Colours indicate focal lineage, error-bars indicate ± 1S.E.M. The fitted blue line is the output of a linear mixed effects model. For each unique focal lineage – non-self pair, n =3 (three biological replicates, with three technical replicates, and all lineages in full interaction with each other).

Environmental quality affects variation in responses. Under carbon enrichment, within-lineage variation is larger and between-lineage variation more pronounced than in ambient-carbon dioxide conditions, although the fold changes in growth rate induced by the presence of a conspecific are lower (see supporting tables 1-2 and 5-6 for statistics). This is especially pronounced when lineages are grown along-side the GFP transformed strain, where, unlike under ambient pCO_2_, a few lineages did not increase their growth rates in response to the GFP strain. Elevated pCO_2_ lineages are already growing faster than their ambient-evolved counterparts, even after accounting for growth under long-term pCO_2_ selection being reduced compared to the short-term effects of elevated pCO_2_ (see ^5^).

### The relationship between monoculture and mixed culture growth is described by a regression to the mean and is consistent with changing energy allocated to growth

Regardless of selection history, intraspecific variation in growth decreases in mixed culture relative to monoculture and shows a pattern consistent with regression to the mean (see also supporting information Fig. 1). This is consistent with there being a range of viable lineage growth rates, bounded by the minimum cell division rate needed for lineage persistence at the lower end, and the cell division rate when the maximum energy is allocated to it on the upper end. In a representative sample of a multi-lineage population, the range of monoculture growth rates under ideal conditions would then estimate the range of growth rates in the multi-lineage population. When a focal lineage responds to the presence of a non-self lineage, the probable direction of their growth rate response is dictated by their monoculture growth rate. Lineages with extremely high monoculture growth rates cannot increase it more, and responses must involve allocating energy to other traits. In contrast, those growing very slowly cannot allocate less energy to growth, so must use a higher-growth strategy in responses.

The above depends on lineages varying in how much energy they allocate to growth. To test whether lineages vary in photosynthetic energy allocation to growth in monoculture, we measured the relationship between biomass gain and net photosynthesis for all lineages of *Ostreococcus* grown in monoculture. We found that lineages differ in their relationships between net photosynthesis (NP) and biomass production (Fig. 1, Fig. 2 D-F, Fig. 3 D-F). Under ambient pCO_2_ (supporting Fig. 2), all lineages, regardless of treatment or lineage identity used less carbon for growth than they produced via net photosynthesis (i.e. the ratio of NP/growth is always larger than 1). Under elevated pCO_2_ (supporting Fig. 3), most lineages used less carbon for growth than they produced via net photosynthesis, however, the surplus in NP was on average lower than in ambient selected lineages (see supporting Fig. 4), and some lineages grew faster than possible by using carbon from NP alone, indicating that they may have been using carbon from storage, or organic carbon retrieved elsewhere. To test whether the reactions of lineages to non-self conspecifics is consistent with energy reallocation, we calculated the percentage of surplus NP (see Fig 1. D), and compared it to how much their growth rate changed when in mixed culture (Fig. 2 D-F, Fig. D-F). While not necessarily a true mechanistic explanation, the relationship holds strong predictive power: Lineages allocating less carbon to growth in monoculture responded more to cues of non-self lineages (and increased growth rates) than lineages allocating more carbon to growth in monoculture, with % of surplus NP explaining up to 50% of the variation in responsiveness. Under elevated pCO_2_, the relationship between the magnitude and direction of change in growth rate in response to a conspecific, and the amount of surplus C from NP becomes more complex, allowing for multiple strategies of storing and allocating carbon (supporting Fig. 5, supporting tables 8 and 9). These data support the interpretation that between-lineage variation in the relationship between monoculture and mixed culture growth reflects diverse energy allocation strategies, and that diverse energy allocation strategies are particularly pronounced in ameliorated environments.

### The potential role of environmental quality in growth-competition relationships

The existence of multiple nearly-equivalent strategies assumes that growth strategies are determined by trade-offs between allocating energy to fitness-related traits ^19-21^. We suggest that, all else equal (i.e. – with similar genetic and physiological capabilities), the number of available strategies that can occur in a given population of closely related individuals is determined by environmental quality, stability and predictability. Here, we focus on the potential role of environmental quality, and find more variation in growth rate modulation in response to non-self conspecifics in high CO_2_ environments (which have more energy in that they can support higher overall population growth rates and represent higher quality environments here) than in ambient CO_2_ environments (which represent lower quality environments here). (supporting Fig. 5) This supports our hypothesis that environmental amelioration not only allows more rapid lineage growth, but could also support more ways for lineages to modulate their growth strategies in response to social milieu.

Our hypothesis on the relationship between environmental quality and the number of possible viable growth strategies leads to the idea that in all but the poorest-quality environments, closely-related lineages can (and should be expected to) vary in how they modulate their growth strategies in response to social milieu within a given abiotic environment. This is also a plausible explanation for the differences between microbial evolution literature that uses extremely poor environments ^22^, and aquatic microbiology literature, which tends to use more moderate environments ^23^, at least with respect to environmental effects on cell division rates.

In poor-quality stable environments, which are commonly used in classical experimental evolution ^22,24^, one strategy will initially be the best, and possibly only viable, strategy in the new environment. This will likely be a strategy involving faster growth due to higher affinity nutrient uptake or tolerance to a toxin or stress, and be associated with high absolute and relative fitness gains. This is in line with experimental findings where the faster growing (locally-adapted) strain wins intraspecific competitions ^25^ or competitions between functional groups ^26^ on average, under abiotic conditions that favour rapid lineage growth. It may be impossible to decrease lineage growth rate (cell division rate) without decreasing overall fitness in poor environments because most microbial evolution experiments use semi-continuous batch culture, which selects for rapid growth, and imposes a minimum population growth rate needed to avoid extinction. However, if populations can adapt enough to poor-quality environments to increase population growth beyond rates needed to avoid extinction by dilution, other strategies evolve, such as the ability to metabolise an alternate food source ^27^.

In contrast, in the moderately stressful or even ameliorated environments used in marine microbial evolution experiments ^23,28-30^, lineage growth rates are high, and risks of extinction by dilution is low. Here, multiple equivalent strategies may have high fitness in the new environment. There is some evidence that maintaining extremely rapid cell division rates in enriched environments can lead to the accumulation of cellular damage and thus select for slower cell division rates to allow damage to be repaired ^31^. It is worth noting that most environments used in marine microbial experiments exploring evolutionary responses to deleterious environmental change do not reduce population growth rates enough to risk extinction during experiments (about 10%) ^32^, though extreme cases of growth reductions of closer to 80% do exist in thermal tolerance experiments ^33^. In contrast, selection environments used in classical microbial evolution experiments are often toxic or low-nutrient enough to cause population extinctions ^34^ or sustained low growth ^35^, though most papers only report changes in relative fitness in mixed culture, and do not report absolute reductions in monoculture growth rates.

Multiple studies, including this one, show that relative lineage growth rates in monoculture is not a consistent predictor of relative lineage growth rates in mixed culture ^4-6,26^ even in stable environments, and under conditions that favor rapid growth. We propose that variation in growth strategies is attributable to lineage-specific reactions to social milieu, and that these reactions are exacerbated in ameliorated environments. Because of this, growth strategies of a focal lineage will depend on their social milieu, as well as the quality of the environment. In addition, we argue that lineage growth is one of many components of fitness, such that many nearly-equivalent energy allocation strategies can exist within populations. Together, these mean that no consistent correlation between monoculture and mixed culture growth should be expected for genetically-diverse populations of phytoplankton, even of a single species. We hypothesised that the number of growth strategies available to a group of closely related lineages depends party on environmental quality, and show data consistent with this hypothesis, where variation in growth modulation in response to the presence of non-self conspecifics is higher in an ameliorated (high CO_2_) environment.

Within-species variation in responses to conspecifics is in line with the emerging pattern that intraspecific variation plays an important role in determining the dynamics and function of phytoplankton populations ^6,14,18^ or functional groups ^7^. Intraspecific variation in responses to biotic cues poses problems with scaling up from monoculture growth rates and functional traits to population composition and function, because the trait values measured in monoculture are unlikely to reflect those in mixed culture for a given lineage. From here, how can we move towards a more nuanced understanding of microbial growth strategies and their relationship with population-level traits such as primary production or nutrient uptake?

We have shown that one useful way to interpret population growth rates in single lineage cultures is as a measure of how much energy is being devoted to making biomass under defined conditions. In this case, the range of growth rates measured in a group of lineages, each grown in monoculture, estimates the upper and lower limits of how much biomass lineages in that population can produce based on maximum or minimum allocations of energy to growth. This assumes that any strategy that we detect in the laboratory was present at high enough frequencies in the wild populations to be sampled in the first place, and is thus not likely to be a low-fitness strategy. One interesting outcome of using an expectation that genetically diverse microbial populations maintain a large number of strategies is that the range of lineage growth rates in monoculture should reflect the range of lineage growth rates possible in a diverse population, though the growth rate of any particular lineage in mixed population cannot be predicted from its growth rate in monoculture. Competition experiments, when paired with monoculture growth experiments, should measure the range of energy reallocation responses possible, and provide clues about the number of strategies in the population. This highlights that the variation in relationships between lineage growth in monoculture and competitive outcomes is not a disagreement between two methods, one of which must be the correct one for estimating lineage growth. Instead, variation in monoculture and mixed culture growth relationships carries important information about the role of traits other than lineage growth rates in determining fitness in diverse microbial populations. It also suggests that the reason growth and lineage traits in monoculture often fail to predict traits such as primary production in mixed culture is due to undersampling population trait distributions. If so, then predictions of population trait distributions could be improved by increasing sampling and the development of higher-throughput lineage trait measurements.

## Conclusions

This study was motivated by a need to understand why so many different relationships between lineage growth rate and lineage competitive ability are observed – why, even under controlled laboratory conditions, competition theory does not reliably predict competitive outcomes in aquatic primary producers. The consequences of making poor predictions about the lineage composition of communities of aquatic primary producers are potentially large; projecting changes to ecologically and biogeochemically important traits such as primary production and nutrient uptake by primary producers depends on scaling up from laboratory studies done on lineages growing in monoculture or in very simplified assemblages. Our data are consistent with a diversity of energy allocation strategies and the modulation of these strategies based on social milieu, and we propose that this provides a single overarching explanation for the diversity of relationships between lineage growth and competitive ability, even within single environments.

One of the goals of global change biology is to project the properties of future populations of aquatic primary producers. Currently, this undertaking is limited by our mechanistic understanding of how the traits of individual lineages and those of populations are linked. Some aspects of intra-specific diversity in growth strategies are simple to accommodate in understanding bulk traits of phytoplankton populations. For example, using trait distributions measured in monoculture will improve estimates of the diversity of strategies possible for a species or functional group and allow *post-hoc* explanations of population properties, such as primary production. In these cases, it would not be possible to map the role of each individual lineage to a population property. Understanding how changes in lineage frequencies affect population properties based on the physiology of single lineages, on the other hand, will require ‘omics approaches where the presence (and eventually frequencies) of particular functions are inferred directly from the genes or gene expression products during growth in diverse populations. In particular, this work highlights first, that population-level predictions that are based on laboratory monoculture studies using one or few lineages (where the distributions of trait values are undersampled) should be interpreted as one sample from a distribution of strategies, rather than as a representative result. Second, we stress that general, mechanistic explanations linking the physiology of individual lineages to population-level traits are vital for using experimental studies to make accurate projections of the composition and properties of future populations.

## Methods

### This is a methods summary. Detailed methods are provided as part of the supporting information Culture of *Ostreococcus* lineages

In this study, we have used six representative lineages from a selection experiment carried out on 16 lineages (described in ^5,7^). Samples were propagated in semi-continuous batch culture during exponential growth, with the inoculum at 100 cells mL^-1^ in 20mL vented tissue culture flasks under the conditions detailed in ^5^.

For the experiments carried out here, we picked at least six lineages that spanned the full magnitude of short-term and evolutionary responses to elevated pCO_2_ (ranging from small responses in lineages isolated near the deep chlorophyll maximum, to the largest responses in the surface lineages).

### Flow cytometry

We used FACS CANTO and DIVA flow cytometres for cell counts (‘event numbers’), as well as green, orange, and red fluorescence. Details on threshold settings and calibration are in ^36^. All analyses were carried out in the R environment *via* the Bioconductor packages.

### Photosynthesis measurements

We measured gross (GP, i.e. photosynthesis rates including respiration) and net (NP, i.e. photosynthesis rates after ‘losses’ to respiration have been subtracted as NP=GP-R) photosynthesis rates in a Clark-type electrode as described in ^5^. Following conversion factors after ^37,38^ and taking into account the size and cellular stoichiometry for the lineages used here (details in ^36^), we converted these measurements from µmol O_2_ per cell and hour to µg carbon produced per µg carbon present as biomass in the sample per hour.

### Indirect and direct co-culture

#### Indirect co-culture using ThinCerts

To test the responses of lineages to the presence from non-self conspecifics, we used ThinCert™ cell culture inserts. An insert is a well with a 0.4µm-membrane base that is suspended into the individual wells of a 12-well plate. It permits extracellular products including nutrients to diffuse but prevents cells from doing so. Here, we used six *Ostreococcus* lineages. The number of lineages was chosen based on a power analysis. For all assays, we inoculated the compartments inside and outside the insert with the same number of cells (100 cells mL^-1^). For the mono-culture ‘control’ conditions, both compartments were inoculated with the same lineage. For the indirect co-culture assays, the lineages were grown in a full factorial setting, with three (evolved) biological replicates and three technical replicates. Samples were distributed so that no one lineage was present solely in the outside or inside compartment in either combination. Growth rates were tracked for each compartment separately.

### Indirect responses to ‘spiked’ media

We used the same six lineages as above, again as a set of three evolved biological replicates and three technical replicates, to test whether we could elicit a response to the perceived presence of non-self conspecifics. To do so, lineages were supplemented with the 0.2µm filtered supernatant (‘spike’) of either the same or a different lineage in a full-factorial design. Growth was tracked as described using a flow cytometer as described above for a period of seven days after the samples had received the spikes.

### Direct co-culture

We co-cultured eight representative *Ostreococcus* lineages with a GFP-transformed Oth95 lineage. For the co-culture experiment, 20 mL of medium were inoculated with 100 cells mL^-1^ of wild-type lineages and GFP populations each. Cell numbers for each population were measured every day for 14-days (two batch-transfers) by flow cytometry. For better comparison with the ThinCert and ‘spike’ experiments (see above), one 12-well plate of GFP lineages was run along-side the experiment in tissue flasks. There was no significant effect of culture in flasks or plates on the relative changes in growth rates or competitive abilities of the GFP- or the wild type lineages.

### Statistical analysis and simulations

All statistical analysis, simulations, and constructions of conceptual figures were carried out in the R environment (final analyses carried out in version 3.5.0, R Core Team (2018). R: A language and environment for statistical computing. R Foundation for Statistical Computing, Vienna, Austria. URL https://www.R-project.org/.). Details on model construction and selection as well as references for all packages used can be found in the supporting information; a summary is provided below. All R code will be made available at the time of manuscript acceptance.

### Flow cytometry data

Flow cytometry data were imported into the R environment through the Bioconductor packages FlowCore (version 1.11.20) and FlowViz (version 1.44). Thresholds on the size (FSC-H) and chlorophyll fluorescence channels (FL3-H) were set within R so that debris and dead cells were not included in any counts. Growth rate of any focal lineage regardless of the specifics of the experimental set-up was calculated as

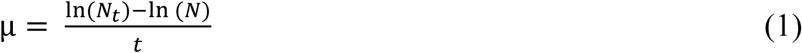

with N_t_ number of cells after a time period t, N number of cells at inoculation, and t the time passed in days.

### Responses to direct (GFP), indirect (ThinCert), or perceived (Spike) presence of non-self conspecifics

Responses to direct (GFP), indirect (ThinCert), or perceived (Spike) presence of other lineages were calculated as the fold-change of

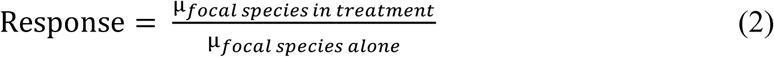

A response of 1 indicates no difference in population growth between monoculture and mixed culture. Values greater than 1 indicates increased growth rates in the direct, indirect, or perceived presence treatments and values less than 1 indicates reduced growth rates.

The shape of the response calculated in (2) as a function of growth in monoculture results in an approximately L-shaped function, and we examined this relationship through a series of non-linear mixed models within the R packages nlme (version 3.1-137) for model fitting, MuMIn (version 1.42.1) for comparison of models and lsmeans (version 2.30-0) for post-hoc tests. For all three cases of mixed culture *vs* monoculture, we defined the same exponential decay function with a variable for slope *a* and intercept *b*. The non-linear mixed effects model was then fitted to that function with the fold change in growth rate as the response variable, and growth in mono-culture as the explaining variable, with fixed effects *a* and *b*. Lineage identity was fitted as a random effect. The global model assumes an effect on both *a* and *b*. Subsequent models were simplified to include an effect on either parameter alone. Models were compared based on their AICc values, and the model with the smallest AICc value chosen for further analysis.

### Carbon allocation and reactiveness to non-self conspecifics

We first plotted the amount of biomass (in µg carbon) produced per hour as a function of net photosynthesis (in µg carbon per µg carbon and hour) (Fig. S1), and analysed the relationship between the two through a linear mixed effects model to account for evolved samples being related to each other in ways that we cannot further disentangle. There, lineage nested within biological replicate was fitted as a random effect, and net photosynthesis was fitted to explain variation in biomass production.

For each biological replicate of each lineage, we then calculated the ratio of net photosynthesis in units carbon to growth in units carbon.

In the next step, we examined the reactiveness of lineages (i.e. the result of (2)) as a function of the ratio between biomass production and NP. For each scenario (i.e. ThinCert, spike, or mixed culture with the GFP strain), we fitted a separate linear mixed model as above.

### Models for conceptual graph

For supporting Fig. 1, we tested whether we would expect to see an L –shaped relationship between growth in monoculture and growth in co-culture as a result of regression to a mean. Here, we established two normal distributions with growth rates ranging from 0.45 (day^-1^) to 1.1 (day^-1^), which are values commonly observed for *Ostreococcus* under full nutrients, saturating light levels, and ambient pCO_2_ ^7,39^. We assume one of these normal distributions to represent growth in monoculture, and the other, growth in mixed culture. From these normal distributions, we randomly draw 1000 samples and their associated growth rates, and calculate (2) as above. As this is essentially a regression to a mean, the resulting relationship is L-shaped as in our experimental data. Figure 1 in the supporting information was created within ggplot2 in R. The figure is for conceptualisation only.

## Acknowledgements

SC was supported by a Royal Society (UK) University Research Fellowship and an European Research Council (ERC) starter grant under the European Community’s Seventh Framework Program. CES was supported by a Scottish Universities Life Science Alliance Scholarship during this work.

## Conflict of Interest

The authors declare no conflict of interest.

