## Supporting Information for "Diverse strategies link growth rate and competitive ability in phytoplankton responses to changes in CO_2_ levels"

**Content**:

#### Detailed Methods

#### Tables:

**Supplementary Table 1 |** Model selection for non-linear mixed effect model fits used in Figure 2 A-C
**Supplementary Table 2 |** Model output for non-linear mixed effect model fits used in Figure 2 A-C
**Supplementary Table 3 |** Model selection for linear mixed effect model used in Figure 2 D-F
**Supplementary Table 4 |** Model output for linear mixed effect model used in Figure 2 D-F

**Supplementary Table 5 |**: Model selection for non-linear mixed effect model fits used in Figure 3 A-C
**Supplementary Table 6 |** Model output for non-linear mixed effect model fits used in Figure 3 A-C
**Supplementary Table 7 |** Model selection for linear mixed effect model used in Figure 3 D-F
**Supplementary Table 8 |** Model output for linear mixed effect model used in Figure 3 D-F

**Supplementary Table 9 |**: Model selection comparing allocation strategies (see Figure 4 in main text)
**Supplementary Table 10 |**: Model output comparing allocation strategies (see Figure 4 in main text)

#### Figures

**Supplementary Figure 1 |** A modelled population where growth rates are randomly drawn from normal distributions.

**Supplementary Figure 2 |** Biomass production in µgC per hour as a function of net photosynthesis (NP) in µgC produced per µgC in the sample per hour for samples evolved at ambient $CO_2$ levels (400ppm)

**Supplementary Figure 3 |** Biomass production in µgC per hour as a function of net photosynthesis (NP) in µgC produced per µgC in the sample per hour for samples evolved at elevated $CO_2$ levels (1000ppm)

**Supplementary Figure 4 |** Percentage of % µgC from net photosynthesis not directly allocated to growth in samples evolved at 400 and 1000ppm $CO_2$

**Supplementary Figure 5 |** Relative importance of carbon allocation strategies in ambient and elevated $pCO_2$ selected lineages in the indirect (ThinCert), perceived (Spike) or direct presence of lineages from the same species complex

#### SI references and R package references

### DETAILED METHODS

#### Culture of *Ostreococcus* lineages

In this study, we have used lineages from a selection experiment described in (Schaum *et al.* 2012; Schaum & Collins 2014). Briefly, 16 lineages of *Ostreococcus* with distinct temperature, light, and salinity preferences were evolved under stable and fluctuating, ambient, and elevated $pCO_2$ for approximately 400 generations. Lineages used for the study presented here had been evolved under stable ambient and stable high $pCO_2$.The associated details for carbon chemistry can be found in Schaum and Collins (2014) Samples were propagated in semi-continuous batch culture during exponential growth, with the inoculum at 100 cells $mL^{-1}$ in 20mL vented tissue culture flasks.

Throughout the selection experiments and all assays, *Ostreococcus* were cultured in 0.2µm-filter-sterilised artificial seawater enriched with vitamins and Keller Medium (Keller *et al.* 1987). Light intensity was set to ~170µmol quanta $m^{-2}$ $s^{-1}$ using LEE Filters foil ('ocean blue'), salinity 32 and temperature 18ºC.

For the experiments carried out here, we picked at least six lineages that spanned the full magnitude of short-term and evolutionary responses to elevated $pCO_2$(ranging from small responses in lineages isolated near the deep chlorophyll maximum, to the largest responses in the surface lineages).

#### Flow cytometry

We used FACS CANTO and DIVA flow cytometres to determine cell counts ('event numbers'), orange fluorescence and red fluorescence (FL2 and FL3 respectively, which together span the photo-pigments most prevalent in *Ostreococcus*) in all lineages, and green fluorescence (FL1) in a green fluorescent protein (GFP)-modified *Ostreococcus* strain. Details on threshold settings and calibration details can be found in (Schaum *et al.* 2015). Data were exported in their raw format and all analyses carried out in the R environment *via* the Bioconductor packages (see below for details on statistics). While the bacterial co-inhabitants were not the main focus of this study, their numbers were regularly assessed via SYBR Gold staining of the cultures (detection via FL1). Bacteria population sizes were stable throughout the experiment.

#### Photosynthesis measurements

We measured gross (GP, i.e. photosynthesis rates including respiration) and net (NP, i.e. photosynthesis rates after 'losses' to respiration have been subtracted as NP=GP-R) photosynthesis rates in a Clark-type electrode as described in (Schaum & Collins 2014). Briefly, cultures were centrifuged to be brought to a density of $10^6$ cells $mL^{-1}$ to maximise the signal to noise ratio in the electrode, and incubated at 18ºC in the dark for 10 minutes. Samples were measured under constant stirring and at their growth temperature (18ºC). Photosynthesis rates were calculated as the slope of the increase in oxygen concertation at 300µmol quanta $m^{-2}$ $s^{-1}$. These light intensities are higher than those used in the incubators to account for self-shading at cell counts necessary to achieve good signal-to-noise ratios in the oxygen electrode. Respiration was measured as the slope of the decrease in oxygen throughout five minutes in the dark. Following conversion factors after (Falkowski *et al.* 1985; Montagnes *et al.* 1994) and taking into account the cellular stoichiometry for the lineages used here (stoichiometry published in (Schaum *et al.* 2015)), we then converted these measurements from µmol $O_2$ per cell and hour to µg carbon produced per µg carbon present as biomass in the sample per hour. To account for respiration of the bacterial co-inhabitants, a 1.0µm filtered fraction of samples was regularly tested on the oxygen electrode. The slope of oxygen consumption in the bacterial fraction was negligible and indiscernible from drift within the electrode. It was therefore not included in the analysis.

#### Indirect and direct co-culture

#### Indirect co-culture using ThinCerts

To test the responses of lineages to the presence from non-self conspecifics, we used ThinCert™ cell culture inserts. An insert is a well with a 0.4µm-membrane base that is suspended into the individual wells of a 12-well plate. It permits extracellular products including nutrients to move passively between insert and well but prevents cells from doing so (see also Figure 1 in the main . Here, we used six representative *Ostreococcus* lineages. The number of lineages was chosen based on a power analysis. For all assays, we inoculated the compartments inside and outside the insert so that they contained the same number of cells to begin with (100 cells $mL^{-1}$). For the mono-culture 'control' conditions, the inside and outside compartment were inoculated with the same lineage. For these indirect co-culture assays, the lineages were grown in a full factorial setting, with three (evolved) biological replicates and three technical replicates. Samples were distributed so that no one lineage was present solely in the outside or inside compartment in either combination. Growth rates were tracked for each compartment separately.

#### Indirect responses to 'spiked' media

We used the same six lineages, again as a set of three evolved biological replicates and three technical replicates, to test whether we could elicit a response to the perceived presence of non-self conspecifics. To do so, lineages were supplemented with the 0.2µm filtered supernatant ('a spike') of either the same or a different lineage in a full-factorial design. Growth was tracked as described using a flow cytometer as described above for a period of seven days after the samples had received the spikes. (see also Figure 1B of main document)

#### Direct co-culture

Finally, we co-cultured eight representative *Ostreococcus* lineages with a GFP transformed Oth95 lineage (Oth95 had proven to be the most amenable to genomic transformation, see also (van Ooijen *et al.* 2012) ). A higher number of lineages was used here to account for the fact that here was only one GFP transformed lineage. For the competition experiment, 20 mL of medium were inoculated with 100 cells $mL^{-1}$of wild-type lineages and GFP populations each. Cell numbers for each population were recorded every day for a 14-day period (two batch-transfers) on a flow-cytometer as described above. For better comparison with the ThinCert and 'spike' experiments (see above), one 12-well plate of GFP lineages was run along-side the experiment in tissue flasks. There was no significant effect of culture in flasks or plates on the relative changes in growth rates or competitive abilities of the GFP- or the wild type lineages. (See also Figure 1C in main document)

#### Statistical analysis and simulations

All statistical analysis, simulations, and constrictions of conceptual figures were carried out in the R environment (final analyses carried out in version 3.5.0, R Core Team (2018). R: A language and environment for statistical computing. R Foundation for Statistical Computing, Vienna, Austria. URL https://www.R-project.org/.). All R code is available upon request. See below for details on the packages used.

#### Statistical analysis
#### Flow cytometry data

Flow cytometry data were imported into the R environment through the Bioconductor packages FlowCore (version 1.11.20) and FlowViz (version 1.44). Thresholds on the size (FSC-H) and chlorophyll fluorescence channels (FL3-H) were set within R so that debris and dead cells were not included in any counts. Each event logged with the correct size and chlorophyll fluorescence then represents one *Ostreococcus* cell and allows us to track changes in cell number. As we are not concerned with the shape of the growth curve, growth rate of any focal lineage regardless of the specifics of the experimental set-up was calculated simply as

$$\mu = \frac{\ln(N_t) - \ln(N)}{t} \qquad (1)$$

with $N_t$ number of cells after a time period t, N number of cells at inoculation, and t the time passed in days.

#### Responses to direct (GFP), indirect (ThinCert), or perceived (Spike) presence of non-self conspecifics

Responses to direct (GFP), indirect (ThinCert), or perceived (Spike) presence of other lineages were calculated as the fold-change of

$$\text{Response} = \frac{\mu_{focal\ species\ in\ treatment}}{\mu_{focal\ species\ alone}} \qquad (2)$$

A response of 1 indicates no difference in population growth rate between growth in monoculture and growth in co-culture. Values greater than 1 indicates increased growth rates in the direct, indirect, or perceived presence treatments and values less than 1 indicates reduced growth rates.
The shape of the response calculated in (2) as a function of growth in monoculture results in an approximately L-shaped function, and we examined this relationship through a series of non-linear mixed models within the R packages nlme (version 3.1-137) for model fitting, MuMIn (version 1.42.1) for comparison of models and lsmeans (version 2.30-0) for post-hoc tests. For all three cases of mixed culture *vs* monoculture, we defined the same exponential decay function with a variable for slope *a* and intercept *b*. The non-linear mixed effects model was then fitted to that function with the fold change in growth rate as the response variable, and growth in mono-culture as the explaining variable, with fixed effects *a* and *b*. Lineage identity was fitted as a random effect. The global model assumes an effect on both *a* and *b*. Subsequent models were simplified to include an effect on either parameter alone. Models were compared based on their AICc values, and the model with the smallest AICc value chosen for further analysis. In all cases, the model with and effect on *a* and *b* in full interaction was the best model with ΔAICc >>5 (read: the L-shaped relationship between growth in monoculture and growth in mixed culture was significant, and the lineages significantly different in how strongly they reacted to the presence of a conspecific too).

#### Carbon allocation and reactiveness to non-self conspecifics

We first plotted the amount of biomass (in µg carbon) produced per hour as a function of net photosynthesis (in µg carbon per µg carbon and hour) (Figure S1), and analysed the relationship between the two through a linear mixed effects model to account for evolved samples being related to each other in ways that we cannot further detangle. There, lineage nested within biological replicate was fitted as a random effect, and net photosynthesis was fitted to explain variation in biomass production.
For each biological replicate of each lineage, we then calculated the ratio of net photosynthesis in units carbon to growth in units carbon. Values lower than 1 indicate that more carbon is used than is being produced at the time of measurements – this would be the case in cells thriving off storage molecules such as starches and lipids (López-Urrutia *et al.* 2006). For values larger than 1 – the more usual scenario (Halsey & Jones 2015) - it is likely that not all carbon is directly allocated to growth. Values that are only slightly larger than 1 indicate that most carbon is channelled to growth, but some is allocated (or exuded, e.g. (Passow 2002)) elsewhere, and values much larger than 1 indicate that *most* carbon is allocated elsewhere.

In the next step, we examined the reactiveness of lineages (i.e. the result of (2)) as a function of the ratio between biomass production and NP. For each scenario (i.e. ThinCert, spike, or mixed culture with the GFP strain), we fitted a separate linear mixed model as above.

#### Models for schematics and conceptual graphs

For the conceptual figures (Supporting Information Figure 1), we first tested whether we would expect to see an L –shaped relationship between growth in monoculture and growth in co-culture as a result of regression to a mean. Here, we established two normal distributions with growth rates ranging from 0.45 ($day^{-1}$) to 1.1 ($day^{-1}$), which are values commonly observed in *Ostreococcus* under full nutrients, saturating light levels, and ambient $pCO_2$ (Rodríguez *et al.* 2005; Schaum *et al.* 2012). We assume one of these normal distributions to represent growth in mono culture, and the other, growth in mixed culture. From these normal distributions, we randomly draw 1000 samples and their associated growth rates, and calculate (2) as above. As this is essentially a regression to a mean, the resulting relationship is L-shaped as in our experimental data.

#### TABLES

(tables were built directly from model output via the prettify command in the papeR package, version 1.04)

**Supplementary Table 1 | Model selection for non-linear mixed effect model fits used in Figure 2 A and B (ambient $CO_2$ data)** We analysed the shape of the relationship between growth alone and the ratio of growth in direct, indirect, or perceived co-culture to growth alone using a non-linear mixed effects model that fits an exponential decay function with an intercept 'a' and a slope 'b'. We then tested whether there the presence of the second species has an impact on either of these parameters. Model selection was carried out by fist fitting the most complex model to the data and then simplifying the random and fixed effects until all fixed effect coefficients were significant at $p < 0.05$. Both parameters were significant, therefor the full model fit was updated using REML. Df is for degrees of freedom. Models were compared via the small sample-size corrected Akaike Information Criterion (AIC).

**A ThinCert Data**

Remove treatment effect on intercept a

| Details | Model number | df | AIC | logLik | Test | L.Ratio | p-value |
|---|---|---|---|---|---|---|---|
| **full model** | 1 | 15 | **926.96** | -448.48 | | | |
| a removed | 2 | 10 | 1013.94 | -496.97 | 1 vs 2 | **96.98** | **<0.0001** |

Remove treatment effect on slope b

| Details | Model number | df | AIC | logLik | Test | L.Ratio | p-value |
|---|---|---|---|---|---|---|---|
| **full model** | 1 | 15 | **926.96** | -448.48 | | | |
| b removed | 3 | 10 | 930.97 | -455.48 | 1 vs 3 | **14.01** | **< 0.01** |

**B – Spike Data**

Remove treatment effect on intercept a

| Details | Model number | df | AIC | logLik | Test | L.Ratio | p-value |
|---|---|---|---|---|---|---|---|
| **full model** | 1 | 15 | **-1034.34** | 532.17 | | | |
| a removed | 2 | 10 | -912.92 | 466.46 | 1 vs 2 | **131.43** | **<0.0001** |

Remove treatment effect on slope b

| Details | Model number | df | AIC | logLik | Test | L.Ratio | p-value |
|---|---|---|---|---|---|---|---|
| **full model** | 1 | 15 | -1034.34 | 532.17 | | | |
| b removed | 2 | 10 | -911.56 | 465.78 | 1 vs 3 | **132.79** | **<0.0001** |

**Supplementary Table 2| Model output for non-linear mixed effect model fits used in Figure 2 A-C (ambient $CO_2$ data).** Parameter estimates are denoted as 'value' and std.error is for 1 S.E.M. Df are for degrees of freedom.

**A Thincerts**

Fixed effects(a +b ~1+lineage2)

| Curve parameter | lineage | Value | Std.Error | DF | t-value | p-value |
|---|---|---|---|---|---|---|
| a | oth95 | 4.14 | 0.74 | 559 | 5.63 | **<0.0001** |
| a | rcc1107 | 5.96 | 0.77 | 559 | 2.35 | **<0.01** |
| a | rcc1108 | 2.45 | 0.74 | 559 | -2.29 | **<0.01** |
| a | rcc1558 | 4.32 | 0.65 | 559 | 0.28 | 0.78 |
| a | rcc410 | 5.99 | 0.68 | 559 | 2.71 | **<0.01** |
| a | rcc810 | 5.59 | 0.78 | 559 | 1.86 | **<0.05** |
| b | oth95 | -3.03 | 0.49 | 559 | -6.15 | **<0.0001** |
| b | rcc1107 | -3.08 | 0.52 | 559 | -0.08 | 0.93 |
| b | rcc1108 | -2.37 | 0.52 | 559 | 1.26 | 0.21 |
| b | rcc1558 | -2.84 | 0.50 | 559 | 0.38 | 0.71 |
| b | rcc410 | -2.88 | 0.50 | 559 | 0.30 | 0.77 |
| b | rcc810 | -3.36 | 0.52 | 559 | -0.63 | 0.53 |

**B Spikes**

Fixed effects(a +b ~1+lineage2)

| Curve parameter | lineage | Value | Std.Error | DF | t-value | p-value |
|---|---|---|---|---|---|---|
| a | oth95 | 3.61 | 0.18 | 253 | 19.93 | **<0.0001** |
| a | rcc1108 | 3.96 | 0.35 | 253 | 1.00 | 0.32 |
| a | rcc1558 | 2.89 | 0.19 | 253 | -3.77 | **<0.001** |
| a | rcc343 | 3.90 | 0.22 | 253 | 1.31 | 0.19 |
| a | rcc410 | 2.05 | 0.19 | 253 | -8.33 | **<0.0001** |
| a | rcc809 | 2.31 | 0.19 | 253 | -6.88 | **<0.0001** |
| b | oth95 | -1.67 | 0.07 | 253 | -24.08 | **<0.0001** |
| b | rcc1108 | -1.74 | 0.11 | 253 | -0.61 | 0.54 |
| b | rcc1558 | -1.31 | 0.07 | 253 | 4.89 | **<0.0001** |
| b | rcc343 | -1.73 | 0.07 | 253 | -0.69 | 0.49 |
| b | rcc410 | -0.91 | 0.08 | 253 | 9.10 | **<0.0001** |
| b | rcc809 | -1.07 | 0.08 | 253 | 7.38 | **<0.0001** |

**C GFP**

Fixed effects(a +b ~1)

| | Value | Std.Error | DF | t-value | p-value |
|---|---|---|---|---|---|
| a | 9.73 | 0.52 | 39 | 18.58 | **<0.0001** |
| b | -1.70 | 0.09 | 39 | -18.88 | **<0.0001** |

**Supplementary Table 3 | Model selection for linear mixed effect model fits used in Figure 2 D-F (ambient $CO_2$ data)** We analysed the relationship between carbon produced (through net photosynthesis) allocated to growth and the ratio of growth in direct, indirect, or perceived co-culture to growth alone using linear mixed effects model. Fitted slopes and intercept as used in Figure 2D -F were derived from the fixed effects. Model selection was carried out by fist fitting the most complex model with full interactions to the data. delta AICc is the difference in AICc score relative to the model with the lowest value (most parsimonious model) and Weight is the relative support for the model. The best fitting models were selected as those returning the lowest AICc score and the highest AICc weight and are highlighted in bold. The best model included lineage effects and effects on relative carbon allocation on growth rate μ, but not in interaction. As delta AICc in the best model is >2, no model averaging was carried out. The best model was then refit with REML.

**D Thincerts model selection**
**Model formula fixed = growthrate.mu ~ surplus_percent_PS* Lineage, random = ~ 1|biorep**

| # | Intercept | Lineage | percentage | Lineage:percentage | df | logLik | AICc | delta | weight |
|---|---|---|---|---|---|---|---|---|---|
| **4** | **0.47** | **+** | **0.01** | **NA** | **9** | **15.55** | **9.41** | **0.00** | **0.70** |
| 3 | 0.11 | NA | 0.02 | NA | 4 | -0.20 | 11.48 | 2.07 | 0.25 |
| 2 | 1.09 | + | NA | NA | 8 | 8.61 | 14.77 | 5.36 | 0.05 |
| 1 | 1.31 | NA | NA | NA | 3 | -7.39 | 22.50 | 13.09 | 0.00 |
| 8 | -0.04 | + | 0.03 | + | 14 | 21.47 | 125.06 | 115.65 | 0.00 |

**E Spikes model selection**
**Model formula fixed = growthrate.mu ~ surplus_percent_PS* Lineage, random = ~ 1|biorep**

| | Intercept | Lineage | percentage | Lineage:percentage | df | logLik | AICc | delta | weight |
|---|---|---|---|---|---|---|---|---|---|
| **3** | **0.83** | **NA** | **0.00** | **NA** | **4.00** | **26.41** | **-41.74** | **0.00** | **0.98** |
| 1 | 1.01 | NA | NA | NA | 3.00 | 20.65 | -33.59 | 8.15 | 0.02 |
| 2 | 1.01 | + | NA | NA | 8.00 | 31.44 | -30.87 | 10.87 | 0.00 |
| 4 | 0.90 | + | 0.00 | NA | 9.00 | 33.79 | -27.07 | 14.66 | 0.00 |
| 8 | 0.93 | + | 0.00 | + | 14.00 | 35.19 | 97.62 | 139.36 | 0.00 |

**F GFP model selection**
**Model formula fixed = growthrate.mu ~ surplus_percent_PS* Lineage, random = ~ 1|biorep**

| | Intercept | Lineage | percentage | Lineage:percentage | df | logLik | AICc | delta | weight |
|---|---|---|---|---|---|---|---|---|---|
| **4** | **2.88** | **+** | **0.03** | **NA** | **10.00** | **-1.26** | **44.52** | **0.00** | **0.95** |
| 2 | 4.15 | + | NA | NA | 9.00 | -8.10 | 50.57 | 6.05 | 0.05 |
| 3 | 1.72 | NA | 0.02 | NA | 4.00 | -25.81 | 62.12 | 17.60 | 0.00 |
| 1 | 2.76 | NA | NA | NA | 3.00 | -27.43 | 62.27 | 17.74 | 0.00 |
| 8 | 3.82 | + | 0.01 | + | 16.00 | 15.97 | 136.06 | 91.54 | 0.00 |

**Supplementary Table 4 | Model output and parameter estimates for linear mixed effect model fits used in Figure 2 D-F (ambient $CO_2$ data)** CI denotes the (95%) confidence intervals. Std Error is the standard error, df degrees of freedom

**D Coculture in ThinCerts**

| **Parameter** | **Value** | **CI (lower)** | **CI (upper)** | **Std.Error** | **DF** | **t-value** | **p-value** | |
|---|---|---|---|---|---|---|---|---|
| Intercept | 0.47 | 0.07 | 0.87 | 0.18 | 9.00 | 2.64 | 0.03 | * |
| Surplus Percent PS | 0.01 | 0.01 | 0.02 | 0.00 | 9.00 | 3.85 | 0.00 | ** |
| lineage: rcc1107 | 0.10 | -0.15 | 0.35 | 0.11 | 9.00 | 0.89 | 0.40 | |
| lineage: rcc1108 | 0.51 | 0.18 | 0.84 | 0.15 | 9.00 | 3.49 | 0.01 | ** |
| lineage: rcc1558 | 0.02 | -0.27 | 0.32 | 0.13 | 9.00 | 0.19 | 0.86 | |
| lineage: rcc410 | -0.28 | -0.56 | -0.01 | 0.12 | 9.00 | -2.31 | 0.05 | * |
| lineage: rcc810 | -0.09 | -0.34 | 0.15 | 0.11 | 9.00 | -0.87 | 0.41 | |

**E Spikes**

| **Parameter** | **Value** | **CI (lower)** | **CI (upper)** | **Std.Error** | **DF** | **t-value** | **p-value** | |
|---|---|---|---|---|---|---|---|---|
| Intercept | 0.90 | 0.76 | 1.04 | 0.06 | 9.00 | 14.28 | <0.001 | *** |
| Surplus Percent PS | 0.00 | 0.00 | 0.01 | 0.00 | 9.00 | 1.92 | 0.09 | . |
| lineage: rcc1107 | 0.01 | -0.11 | 0.13 | 0.05 | 9.00 | 0.21 | 0.84 | |
| lineage: rcc1108 | -0.08 | -0.18 | 0.03 | 0.05 | 9.00 | -1.69 | 0.12 | |
| lineage: rcc1558 | -0.01 | -0.16 | 0.14 | 0.07 | 9.00 | -0.15 | 0.88 | |
| lineage: rcc410 | -0.10 | -0.20 | -0.01 | 0.04 | 9.00 | -2.39 | 0.04 | * |
| lineage: rcc810 | -0.06 | -0.15 | 0.02 | 0.04 | 9.00 | -1.62 | 0.14 | |

**F GFP**

| **Parameter** | **Value** | **CI (lower)** | **CI (upper)** | **Std.Error** | **DF** | **t-value** | **p-value** | |
|---|---|---|---|---|---|---|---|---|
| Intercept | 2.88 | 1.99 | 3.76 | 0.40 | 11.00 | 7.16 | <0.001 | *** |
| Surplus Percent PS | 0.03 | 0.01 | 0.04 | 0.01 | 11.00 | 3.57 | 0.004 | ** |
| lineage: rcc1107 | -0.36 | -1.01 | 0.29 | 0.30 | 11.00 | -1.21 | 0.253 | |
| lineage: rcc1108 | -2.18 | -2.86 | -1.50 | 0.31 | 11.00 | -7.05 | <0.001 | *** |
| lineage: rcc1558 | -0.89 | -1.49 | -0.29 | 0.27 | 11.00 | -3.27 | 0.007 | ** |
| lineage: rcc410 | -1.06 | -1.67 | -0.45 | 0.28 | 11.00 | -3.81 | 0.003 | ** |
| lineage: rcc810 | -1.66 | -2.25 | -1.07 | 0.27 | 11.00 | -6.23 | <0.001 | *** |

**Supplementary Table 5 | Model selection for non-linear mixed effect model fits used in Figure 3 A and B (elevated $CO_2$ data)** We analysed the shape of the relationship between growth alone and the ratio of growth in direct, indirect, or perceived co-culture to growth alone using a non-linear mixed effects model that fits an exponential decay function with an intercept 'a' and a slope 'b'. We then tested whether there the presence of the second species has an impact on either of these parameters. Model selection was carried out by fist fitting the most complex model to the data and then simplifying the random and fixed effects until all fixed effect coefficients were significant at $p < 0.05$. Both parameters were significant, therefor the full model fit was updated using REML. Df is for degrees of freedom. Models were compared via the small sample-size corrected Akaike Information Criterion (AIC).

**Figure 3A ThinCert Data**

| Details | Model number | df | AIC | logLik | Test | L.Ratio | p-value |
|---|---|---|---|---|---|---|---|
| full model | 1 | 15 | 319.79 | 174.90 | | | |
| parameters removed | 2 | 10 | 369.94 | 194.69 | 1 vs 2 | 36.58 | **<0.0001** |

**B – Spike Data**

| Details | Model number | df | AIC | logLik | Test | L.Ratio | p-value |
|---|---|---|---|---|---|---|---|
| **full model** | 1 | 15 | **-514.26** | 270.13 | | | |
| a removed | 2 | 10 | -512.17 | 265.08 | 1 vs 2 | **10.11** | **<0.01** |

**Supplementary Table 6| Model output for non-linear mixed effect model fits used in Figure 3 A-C (elevated $CO_2$ data).** Parameter estimates are denoted as 'value' and std.error is for 1 S.E.M. Df are for degrees of freedom.

**A Thincerts**

Fixed effects(a +b ~1+lineage2)

| Curve parameter | lineage | Value | Std.Error | DF | t-value | p-value |
|---|---|---|---|---|---|---|
| a | oth95 | 3.45 | 0.19 | 520.00 | 18.18 | **<0.001** |
| a | rcc1107 | 0.33 | 0.26 | 520.00 | 1.26 | 0.21 |
| a | rcc1108 | -0.04 | 0.25 | 520.00 | -0.17 | 0.87 |
| a | rcc1558 | -0.27 | 0.27 | 520.00 | -1.02 | 0.31 |
| a | rcc410 | -0.34 | 0.25 | 520.00 | -1.35 | 0.18 |
| a | rcc810 | -0.18 | 0.26 | 520.00 | -0.70 | 0.49 |
| b | oth95 | -1.30 | 0.07 | 520.00 | -18.40 | **<0.001** |
| b | rcc1107 | -0.14 | 0.10 | 520.00 | -1.40 | 0.16 |
| b | rcc1108 | 0.00 | 0.10 | 520.00 | 0.05 | 0.96 |
| b | rcc1558 | 0.07 | 0.10 | 520.00 | 0.71 | 0.48 |
| b | rcc410 | 0.15 | 0.10 | 520.00 | 1.52 | 0.13 |
| b | rcc810 | 0.07 | 0.10 | 520.00 | 0.68 | 0.50 |

**B Spikes**

Fixed effects list(a ~ 1 + lineage2, b ~ 1)

| Curve parameter | lineage | Value | Std.Error | DF | t-value | p-value |
|---|---|---|---|---|---|---|
| a | | 4.11 | 1.16 | 759.00 | 3.55 | **<0.001** |
| b | oth95 | -1.35 | 0.35 | 759.00 | -3.88 | **<0.001** |
| b | rcc1108 | -0.03 | 0.02 | 759.00 | -1.69 | 0.09 |
| b | rcc1558 | -0.07 | 0.02 | 759.00 | -2.99 | **<0.01** |
| b | rcc343 | -0.03 | 0.02 | 759.00 | -1.41 | 0.16 |
| b | rcc410 | -0.01 | 0.02 | 759.00 | -0.26 | 0.79 |
| b | rcc809 | -0.04 | 0.02 | 759.00 | -1.81 | **<0.05** |

**C GFP**

Fixed effects(a +b ~1)

| | Value | Std.Error | DF | t-value | p-value |
|---|---|---|---|---|---|
| **a** | 6.09 | 0.42 | 63.00 | 14.57 | **<0.0001** |
| **b** | -1.38 | 0.07 | 63.00 | -18.80 | **<0.0001** |

**Supplementary Table 7 | Model selection for linear mixed effect model fits used in Figure 3 D-F ($CO_2$ elevated data)** We analysed the relationship between carbon produced (through net photosynthesis) allocated to growth and the ratio of growth in direct, indirect, or perceived co-culture to growth alone using linear mixed effects model. Fitted slopes and intercept as used in Figure 3D -F were derived from the fixed effects. Model selection was carried out by fist fitting the most complex model with full interactions to the data. delta AICc is the difference in AICc score relative to the model with the lowest value (most parsimonious model) and Weight is the relative support for the model. The best fitting models were selected as those returning the lowest AICc score and the highest AICc weight and are highlighted in bold. The best model included lineage effects and effects on relative carbon allocation on growth rate μ, but not in interaction. Where delta AICc in the best model is >2, no model averaging was carried out. For cases where this was not the case, models were averaged within the MuMIn package until AICc>2.The best model was then refit with REML.

**D Thincerts model selection**

**Model formula fixed = growthrate.mu ~ surplus_percent_PS* Lineage, random = ~ 1|biorep**

| # | Intercept | Lineage | percentage | Lineage:percentage | df | logLik | AICc | delta | weight |
|---|---|---|---|---|---|---|---|---|---|
| **3** | **0.75** | **NA** | **0.01** | **NA** | **4.00** | **-3.39** | **17.85** | **0.00** | **0.92** |
| 1 | 1.10 | NA | NA | NA | 3.00 | -8.41 | 24.54 | 6.69 | 0.03 |
| 2 | 0.90 | + | NA | NA | 8.00 | 3.52 | 24.97 | 7.12 | 0.03 |
| 4 | 0.82 | + | 0.01 | NA | 9.00 | 7.61 | 25.27 | 7.42 | 0.02 |
| 8 | 0.84 | + | 0.00 | + | 14.00 | 30.29 | 107.41 | 89.56 | 0.00 |

**E Spikes model selection**

**Model formula fixed = growthrate.mu ~ surplus_percent_PS* Lineage, random = ~ 1|biorep**

| | Intercept | Lineage | percentage | Lineage:percentage | df | logLik | AICc | delta | weight |
|---|---|---|---|---|---|---|---|---|---|
| **4** | **0.82** | **+** | **0.01** | **NA** | **9.00** | **13.83** | **12.84** | **0.00** | **0.61** |
| **3** | **0.74** | **NA** | **0.01** | **NA** | **4.00** | **-1.48** | **14.04** | **1.21** | **0.33** |
| 1 | 1.05 | NA | NA | NA | 3.00 | -5.02 | 17.76 | 4.93 | 0.05 |
| 2 | 0.96 | + | NA | NA | 8.00 | 5.20 | 21.61 | 8.77 | 0.01 |
| 8 | 0.85 | + | 0.01 | + | 14.00 | 26.92 | 114.17 | 101.33 | 0.00 |

**E GFP model selection**

**Model formula fixed = growthrate.mu ~ surplus_percent_PS* Lineage, random = ~ 1|biorep**

| | Intercept | Lineage | percentage | Lineage:percentage | df | logLik | AICc | delta | weight |
|---|---|---|---|---|---|---|---|---|---|
| **4** | **0.67** | **+** | **0.01** | **NA** | **11.00** | **9.10** | **25.81** | **0.00** | **0.83** |
| **2** | **0.79** | **+** | **NA** | **NA** | **10.00** | **3.94** | **29.05** | **3.24** | **0.17** |
| 3 | 0.91 | NA | 0.02 | NA | 4.00 | -28.13 | 66.37 | 40.56 | 0.00 |
| 1 | 1.89 | NA | NA | NA | 3.00 | -32.52 | 72.25 | 46.44 | 0.00 |
| 8 | 0.75 | + | 0.00 | + | 18.00 | 31.65 | 109.51 | 83.70 | 0.00 |

**Supplementary Table 8| Model output and parameter estimates for linear mixed effect model fits used in Figure 3 D-F (elevated $CO_2$ data)** CI denotes the (95%) confidence intervals. Std Error is the standard error, df degrees of freedom

**D Coculture in ThinCerts**

| Parameter | Value | CI (lower) | CI (upper) | Std.Error | DF | t-value | p-value | |
|---|---|---|---|---|---|---|---|---|
| Intercept | 0.82 | 0.54 | 1.09 | 0.12 | 9.00 | 6.68 | <0.001 | *** |
| Surplus Percent PS | 0.01 | 0.00 | 0.01 | 0.00 | 9.00 | 2.52 | <0.05 | * |
| lineage: rcc1107 | -0.12 | -0.50 | 0.25 | 0.17 | 9.00 | -0.75 | 0.48 | |
| lineage: rcc1108 | -0.33 | -0.76 | 0.10 | 0.19 | 9.00 | -1.74 | 0.12 | |
| lineage: rcc1558 | -0.01 | -0.40 | 0.39 | 0.17 | 9.00 | -0.04 | 0.97 | |
| lineage: rcc410 | 0.34 | -0.10 | 0.79 | 0.20 | 9.00 | 1.74 | 0.12 | |
| lineage: rcc810 | 0.42 | -0.03 | 0.86 | 0.20 | 9.00 | 2.13 | 0.05 | * |

**E Spikes**

| Parameter | Value | CI (lower) | CI (upper) | Std.Error | DF | t-value | p-value | |
|---|---|---|---|---|---|---|---|---|
| Intercept | 0.82 | 0.62 | 1.02 | 0.09 | 9.00 | 9.22 | <0.001 | *** |
| Surplus Percent PS | 0.01 | 0.00 | 0.01 | 0.00 | 9.00 | 4.21 | <0.01 | ** |
| lineage: rcc1107 | -0.49 | -0.82 | -0.16 | 0.14 | 9.00 | -3.38 | <0.01 | ** |
| lineage: rcc1108 | -0.27 | -0.54 | 0.01 | 0.12 | 9.00 | -2.19 | 0.06 | . |
| lineage: rcc1558 | -0.28 | -0.58 | 0.01 | 0.13 | 9.00 | -2.18 | <0.05 | * |
| lineage: rcc410 | 0.18 | -0.17 | 0.54 | 0.16 | 9.00 | 1.17 | 0.27 | |
| lineage: rcc810 | 0.11 | -0.16 | 0.38 | 0.12 | 9.00 | 0.91 | 0.39 | |

**F GFP**

| Parameter | Value | CI (lower) | CI (upper) | Std.Error | DF | t-value | p-value | |
|---|---|---|---|---|---|---|---|---|
| Intercept | 0.67 | 0.39 | 0.95 | 0.13 | 13.00 | 5.24 | <0.001 | *** |
| Surplus Percent PS | 0.01 | 0.00 | 0.01 | 0.00 | 13.00 | 2.91 | <0.05 | * |
| lineage: rcc1107 | 0.47 | 0.11 | 0.84 | 0.17 | 13.00 | 2.81 | <0.05 | * |
| lineage: rcc1108 | -0.09 | -0.48 | 0.30 | 0.18 | 13.00 | -0.49 | 0.64 | |
| lineage: rcc1558 | 1.60 | 1.21 | 1.98 | 0.18 | 13.00 | 8.89 | <0.001 | *** |
| lineage: rcc410 | 1.27 | 0.81 | 1.72 | 0.21 | 13.00 | 5.99 | <0.001 | *** |
| lineage: rcc810 | 0.94 | 0.54 | 1.33 | 0.18 | 13.00 | 5.10 | <0.001 | *** |

**Supplementary Table 9|**: Model selection comparing allocation strategies (see Figure 4 in main text). We analysed how excess carbon produced (through net photosynthesis) is influenced by co-culture method (direct with GFP, indirect with ThinCerts, or perceived with spikes) and $pCO_2$ level (ambient or elevated). Model selection was carried out by fist fitting the most complex model with full interactions to the data. delta AICc is the difference in AICc score relative to the model with the lowest value (most parsimonious model) and Weight is the relative support for the model. The best fitting models were selected as those returning the lowest AICc score and the highest AICc weight and are highlighted in bold. The best model included culture method and $CO_2$ treatment, in full interaction. Where delta AICc in the best model is >2, no model averaging was carried out. For cases where this was not the case, models were averaged within the MuMIn package until AICc>2.The best model was then refit with REML.

**Model formula: fixed =carb.excess ~ ppm*culture.type, random = ~ 1|lineage_biorep**

| | Intercept | ppm | Culture type | Ppm* Culture type | df | logLik | AICc | delta | weight |
|---|---|---|---|---|---|---|---|---|---|
| **8** | **10.87** | **+** | **+** | **+** | **8.00** | **-323.05** | **663.40** | **0.00** | **1.00** |
| 4 | 11.58 | + | + | NA | 6.00 | -333.97 | 680.68 | 17.28 | 0.00 |
| 3 | 9.13 | NA | + | NA | 5.00 | -353.50 | 717.52 | 54.12 | 0.00 |
| 2 | 16.45 | + | NA | NA | 4.00 | -385.42 | 779.19 | 115.79 | 0.00 |
| 1 | 14.01 | NA | NA | NA | 3.00 | -395.87 | 797.95 | 134.55 | 0.00 |

**Supplementary Table 10|**: Model output comparing allocation strategies (see Figure 4 in main text). CI denotes the (95%) confidence intervals. Std Error is the standard error, df degrees of freedom

| **Parameter** | **Value** | **CI (lower)** | **CI (upper)** | **Std.Error** | **DF** | **t-value** | **p-value** | |
|---|---|---|---|---|---|---|---|---|
| (Intercept) | 10.88 | 9.04 | 12.73 | 0.93 | 103.00 | 11.72 | **<0.001** | *** |
| ppm: 1000ppm | -3.33 | -5.68 | -0.99 | 1.18 | 103.00 | -2.82 | **<0.01** | ** |
| react_to: GFP | 13.23 | 10.94 | 15.51 | 1.15 | 103.00 | 11.48 | **<0.001** | *** |
| react_to: SPIKE | -1.39 | -3.81 | 1.02 | 1.22 | 103.00 | -1.14 | 0.26 | |
| ppm1000ppm:react_toGFP | -5.17 | -8.27 | -2.06 | 1.56 | 103.00 | -3.30 | **<0.01** | ** |
| ppm1000ppm:react_toSPIKE | 2.00 | -1.32 | 5.32 | 1.67 | 103.00 | 1.20 | 0.24 | |

#### FIGURES

(all figures were constructed within the ggplot2 package version 3.0.0, using the cowplot add-on)

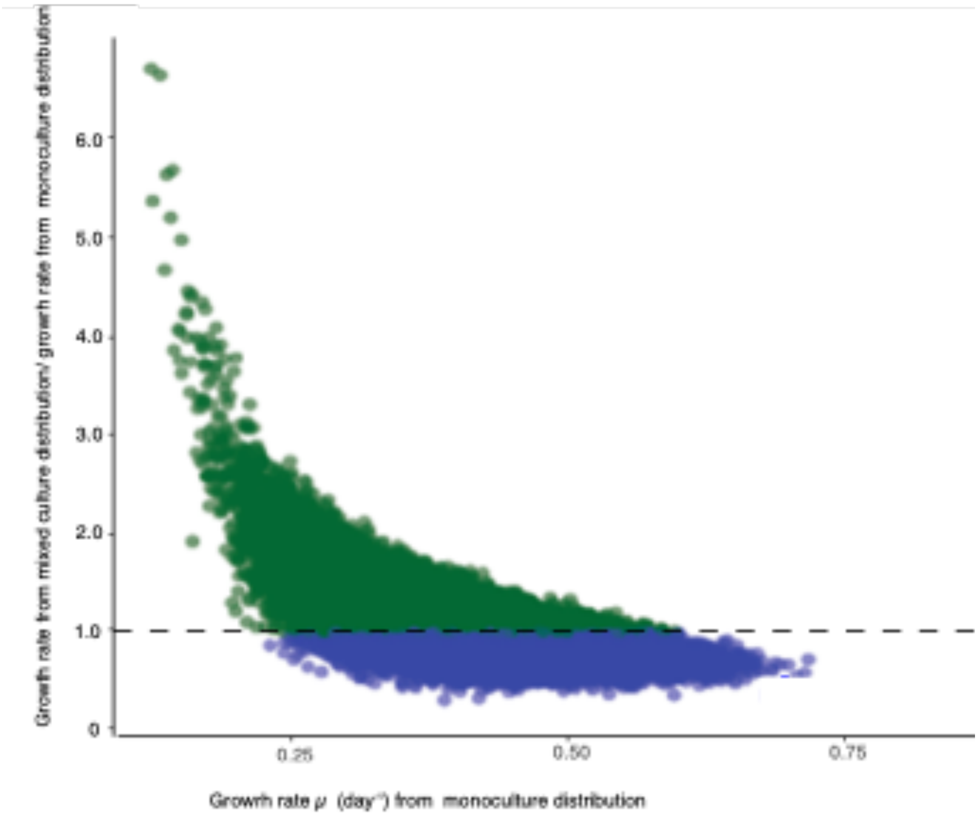


**Supplementary Figure 1 | A modelled population where growth rates are randomly drawn from normal distributions.** The dashed line indicates a fold-change of 1. Values of fold change >1 indicate faster growth in mixed culture than in monoculture. Green denotes fold change values >1, and blue, values <1.

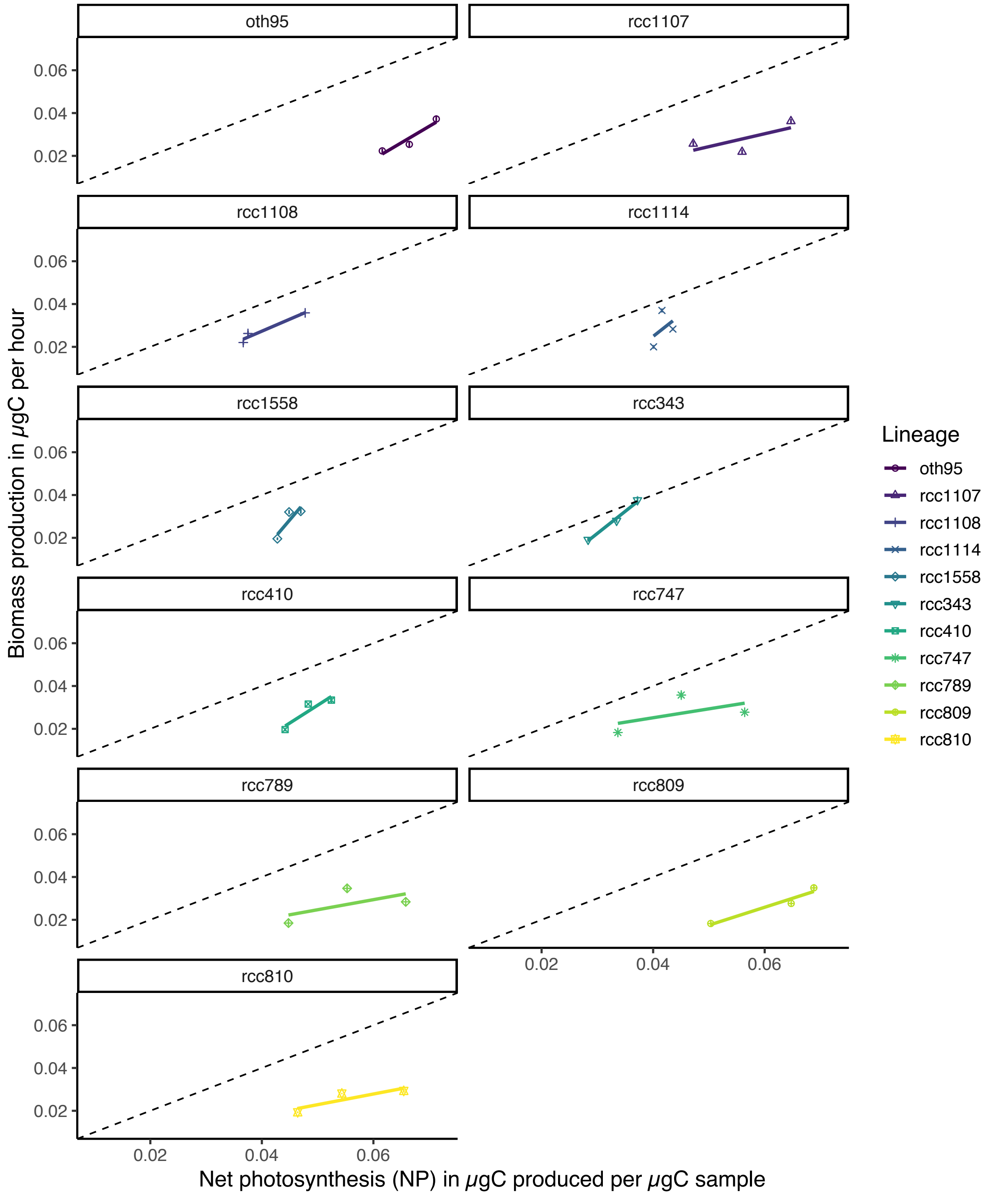


**Supplementary Figure 2 | Biomass production in µgC per hour as a function of Net photosynthesis (NP) in µgC produced per µgC in the sample per hour for samples evolved at ambient $CO_2$ levels (400ppm).** Biomass production and net photosynthesis are positively correlated with each other, such that higher rates of NP yield faster growth. The fitted line is a simple linear model applied on a per lineage basis. In all cases, rates of NP expressed in units carbon are higher (up to two fold) than rates of growth expressed in units carbon, indicating that not all carbon is allocated directly to growth. Colours indicate the different lineages, the dashed line is the 1:1 line where the amount of carbon produced through NP equals the amount of increase in biomass in units carbon. For each lineage n per biological replicate=3. Each biological replicate was measured in triplicates for technical replication.

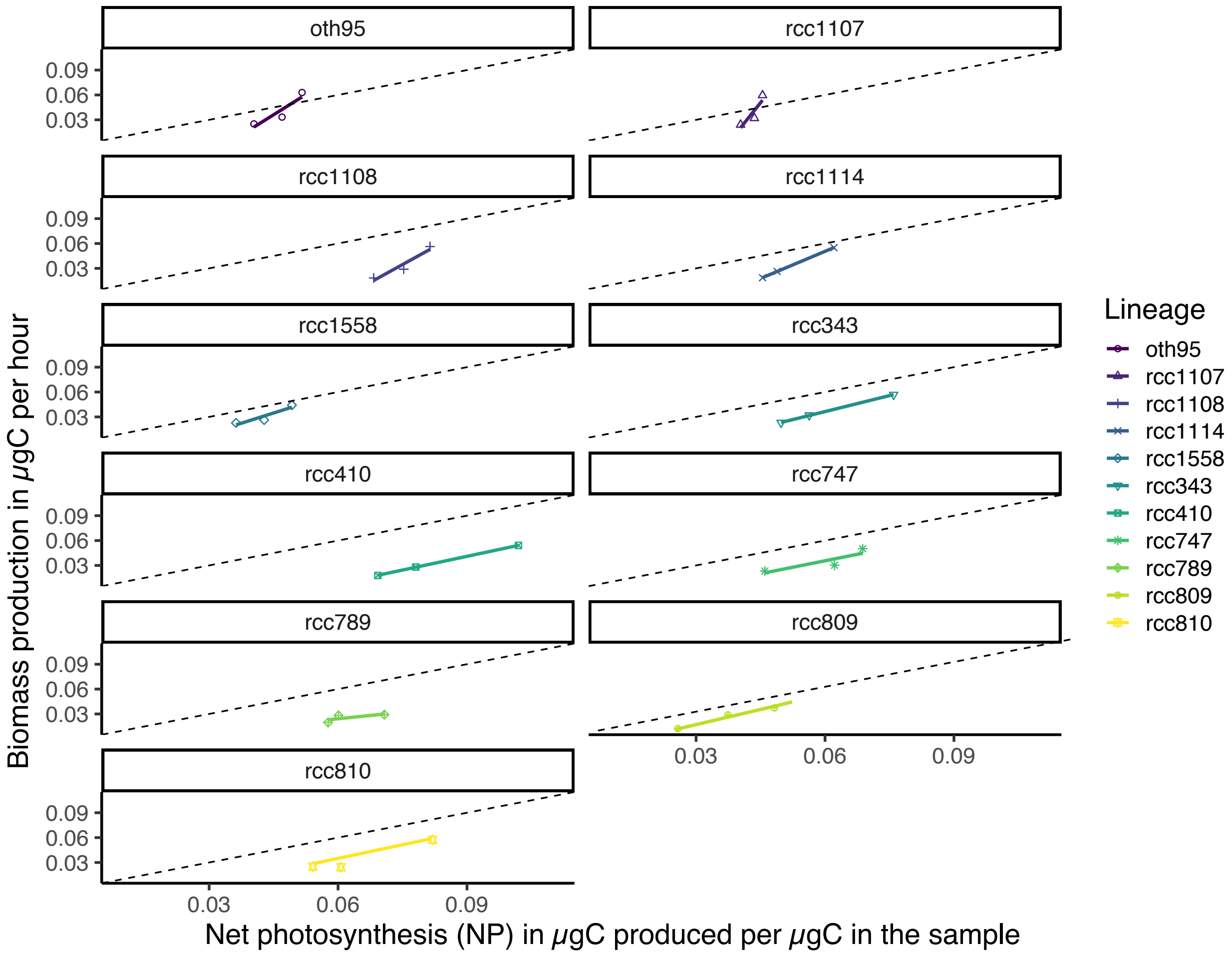


**Supplementary Figure 3 | Biomass production in µgC per hour as a function of Net photosynthesis (NP) in µgC produced per µgC in the sample per hour for samples evolved at ambient $CO_2$ levels (1000ppm).** Biomass production and net photosynthesis are positively correlated with each other, such that higher rates of NP yield faster growth. The fitted line is a simple linear model applied on a per lineage basis. In all cases, rates of NP expressed in units carbon are higher (up to two fold) than rates of growth expressed in units carbon, indicating that not all carbon is allocated directly to growth. Colours indicate the different lineages, the dashed line is the 1:1 line where the amount of carbon produced through NP equals the amount of increase in biomass in units carbon. For each lineage n per biological replicate=3. Each biological replicate was measured in triplicates for technical replication.

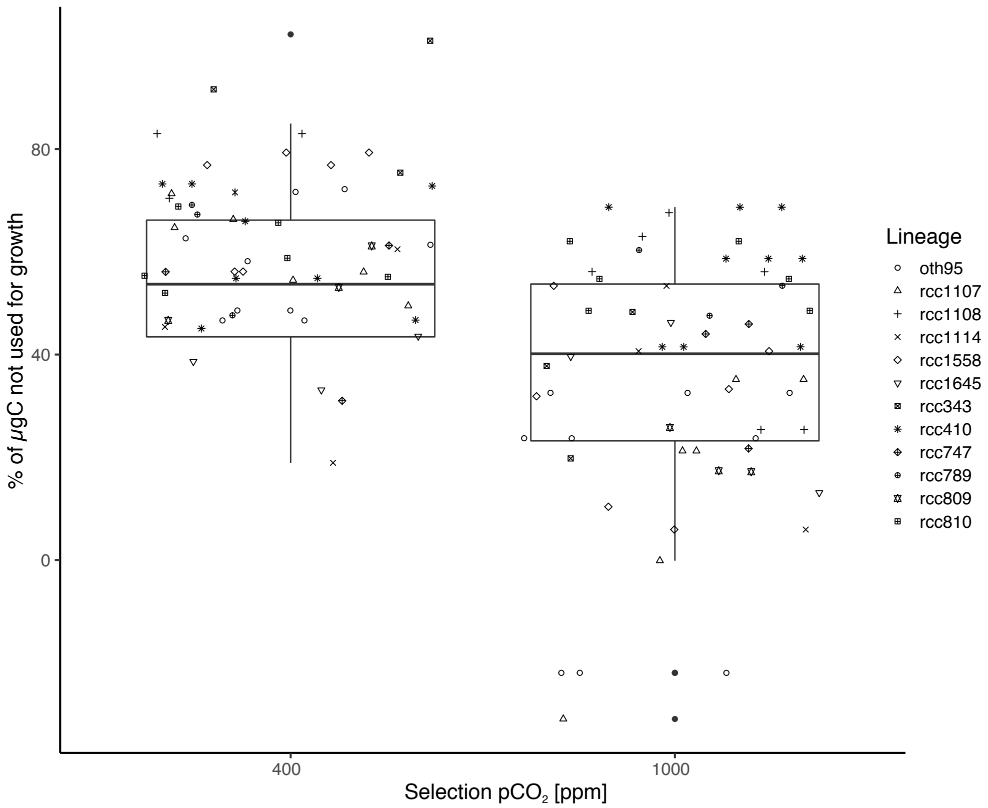


**Supplementary Figure 4 | Selection $pCO_2$ affects how much of carbon produced during net photosynthesis is not directly used for growth.** Under elevated $pCO_2$ significantly less (ANOVA using ppm as a fixed and bioreplicate nested within lineage as a random factor: $F_{1,106} = 12.22$, $p< 0.001$) carbon is directly put into growth of biomass, indicating that under ameliorated circumstances, this 'excess' carbon might be used elsewhere. Symbols denote the lineages used in the experiment. Boxplots are a composite of all co-culture (indirect, perceived, direct) data and are displayed as is standard, with the girdle indicating the median and whiskers extending to the

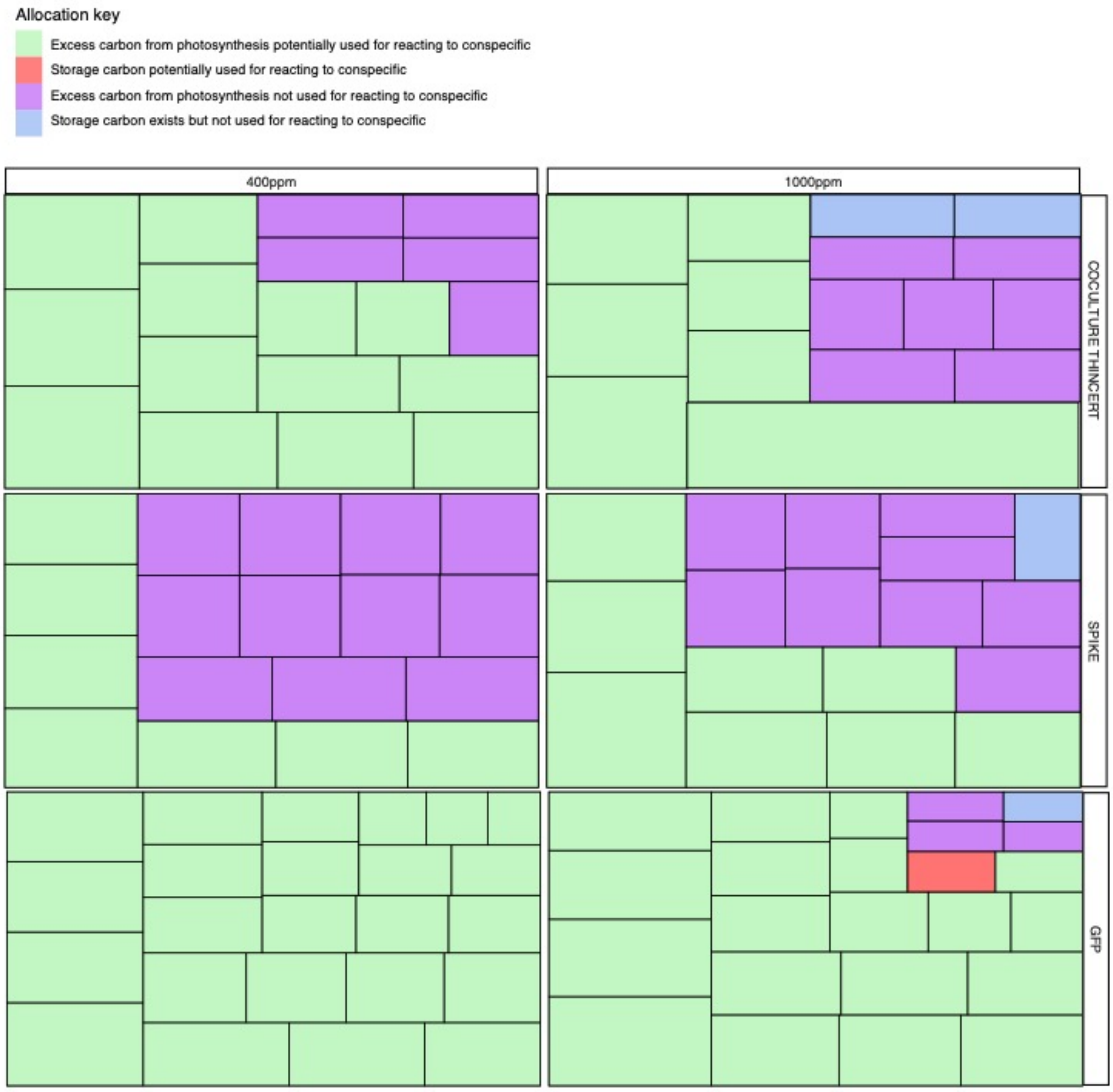


**Supplementary Figure 5 | Relative importance of carbon allocation strategies in ambient and elevated $pCO_2$ selected lineages in the indirect (ThinCert), perceived (Spike) or direct presence of lineages from the same species complex**. Selection $pCO_2$ and culture method influence whether lineages (i) primarily produce excess carbon *and* react readily to conspecifics through increasing their growth rate (green panels), (ii) do not produce excess carbon but must channel storage or other organic carbon to growth *and* react readily to conspecifics through increasing their growth rate (red panels), (iii) primarily produce excess carbon *but do not* react readily to conspecifics through increasing their growth rate (purple panels), (iv) do not produce excess carbon but must channel storage or other organic carbon to growth *but do not* react readily to conspecifics through increasing their growth rate (blue panels). Each sub-panel represents a biological replicate of a focal lineage, and the size of the panel indicates the relative magnitude of the growth response (i.e. a subpanel with a large area indicates high responsiveness to the presence of a conspecific). Regardless of the selection environment, lineages are more likely to simultaneously produce excess carbon and react to conspecifics through faster growth than in mono-culture, the more direct the interaction with a conspecific (compare GFP to spike, for example). However, across all co-culture scenarios, elevated $pCO_2$ selected lineages are more likely to display more diverse strategies than ambient $pCO_2$ selected samples. See supporting tables S9 and S10 for detailed statistics.

#### R package references
#### (order as mentioned in text)

**FlowCore:** B. Ellis, P. Haaland, F. Hahne, N. Le Meur, N. Gopalakrishnan, J. Spidlen and M. Jiang (2018). flowCore: flowCore: Basic structures for flow cytometry data.

**FlowViz:** B. Ellis, R. Gentleman, F. Hahne, N. Le Meur, D. Sarkar and M. Jiang (2018). flowViz: Visualization for flow cytometry. R

**nlme**: Pinheiro J, Bates D, DebRoy S, Sarkar D, R Core Team (2018)._nlme: Linear and Nonlinear Mixed Effects Models_. R package version 3.1-137, <URL:ttps://CRAN.R-project.org/package=nlme>.

**MuMIn**: Kamil Bartoń (2018). MuMIn: Multi-Model Inference. R package version 1.42.1. https://CRAN.R-project.org/package=MuMIn

**lsmeans**: Russell V. Lenth (2016). Least-Squares Means: The R Package lsmeans. Journal of Statistical Software, 69(1), 1-33. doi:10.18637/jss.v069.i01

**prettify**: B. Hofner (2019). papeR: A Toolbox for Writing Pretty Papers and Reports, R package version 1.0-4, https://CRAN.R-project.org/package=papeR.

**ggplot2**: H. Wickham. ggplot2: Elegant Graphics for Data Analysis. Springer-Verlag New York, 2016.

**cowplot**: Claus O. Wilke (2018). cowplot: Streamlined Plot Theme and Plot Annotations for 'ggplot2'. R package version 0.9.3. https://CRAN.R-project.org/package=cowplot
